# Monotherapy efficacy of BBB-permeable small molecule activators of PP2A in glioblastoma

**DOI:** 10.1101/777276

**Authors:** Joni Merisaari, Oxana V. Denisova, Milena Doroszko, Vadim Le Joncour, Patrik Johansson, William P.J. Leenders, David B. Kastrinsky, Nilesh Zaware, Goutham Narla, Pirjo Laakkonen, Sven Nelander, Michael Ohlmeyer, Jukka Westermarck

## Abstract

Glioblastoma (GB) is a fatal disease in which most targeted therapies have clinically failed. However, pharmacological reactivation of tumor suppressors has not been thoroughly studied as yet as a GB therapeutic strategy. Tumor suppressor Protein Phosphatase 2A (PP2A), is inhibited by non-genetic mechanisms in GB, and thus it would be potentially amendable for therapeutic reactivation. Here we demonstrate, that <u>s</u>mall <u>m</u>olecule <u>a</u>ctivators of <u>P</u>P2A (SMAPs), NZ-8-061 and DBK-1154, effectively cross the *in vitro* model of blood-brain barrier (BBB), and *in vivo* partition to mouse brain tissue after oral dosing. *In vitro*, SMAPs exhibit robust cell killing activity against five established GB cell lines, and nine patient-derived primary glioma cell lines. Collectively these cell lines have heterogenous genetic background, kinase inhibitor resistance profile, and stemness properties; and they represent different clinical GB subtypes. Oral dosing of either of the SMAPs significantly reduced growth of infiltrative intracranial GB tumors. DBK-1154, with both higher degree of brain/blood distribution, and more potent *in vitro* activity against all tested GB cell lines, also significantly increased survival of mice bearing orthotopic GB xenografts. In summary, this report presents a proof-of-principle data for BBB-permeable tumor suppressor reactivation therapy for glioblastoma cells of heterogenous molecular background.

## Introduction

One of the hallmarks of GB is dysregulated phosphorylation-dependent pathways, which are drivers of its malignant progression (Dunn *et al.*, 2012). Nevertheless, all tested kinase inhibitors have failed to prolong the overall survival of GB patients in clinical trials (Mooney *et al.*, 2019; Tomiyama and Ichimura, 2019). One of the reasons is that the blood-brain barrier (BBB) prevents the otherwise potentially effective kinase inhibitors from reaching the brain at high enough concentrations (Harder *et al.*, 2018). Another potential reason is non-mutational plasticity induced by kinase inhibitors in GB cells (van den Heuvel *et al.*, 2017). Notably, phosphorylation of GB driver pathways is not only regulated by kinases, but also by phosphatases (Narla *et al.*, 2018; Tomiyama *et al.*, 2019). Amongst them, Protein phosphatase 2A (PP2A) is a serine/threonine phosphatase that regulates multiple oncogenic kinase signaling pathways, as well as apoptotic mechanisms (Perrotti and Neviani, 2013; Kauko and Westermarck, 2018). Interestingly, because PP2A is not genetically inactivated in GB (Kaur *et al.*, 2016), it could be a suitable target for tumor suppressor reactivation therapies in this medically challenging indication.

A series of <u>s</u>mall <u>m</u>olecule <u>a</u>ctivators of <u>P</u>P2A (SMAPs), was recently derived from tricyclic neurological drugs such as Chlorpromazine and Clomipramine (Kastrinsky *et al.*, 2015). Based on photoaffinity labeling studies, radiolytic footprinting, and mutagenesis of the binding site, the target of SMAPs has been identified as the interface of the A and C subunits in the PP2A complex (Sangodkar *et al.*, 2017). The SMAPs have shown efficacy as orally available monotherapy in animal models in solid non-CNS cancer types (Sangodkar *et al.*, 2017; Kauko *et al.*, 2018), but neither their BBB penetration properties, nor potential as CNS cancer drug has not been studied as yet.

## Results and discussion

### Development of blood-brain barrier permeable small molecule reactivators of PP2A

We sought to improve the potency and oral bioavailability of RTC-5 and RTC-30 (Kastrinsky *et al.*, 2015), earlier members of the SMAP series derived from tricyclics, by constraint of the linear spacer moiety between the tricyclic and sulfonamide, resulting in structures exemplified by DBK-1154 and NZ-8-061 (Fig. 1A, and Supplementary figure 1). However, it was unclear whether the polarity introduced to tricyclics by the sulfonamide and hydroxyl moieties (Fig. 1A) would compromise BBB permeability, and eventually CNS availability, of the SMAPs as compared to tricyclics. We therefore started by investigating the *in vitro* BBB passage of NZ-8-061 (a.k.a DT-that has been widely used in cancers outside the CNS (Sangodkar *et al.*, 2017; Kauko *et al.*, 2018; McClinch *et al.*, 2018). Quantified by HPLC-MS/MS, NZ-8-061 was found to cross the artificial BBB (Le Joncour *et al.*, 2019), consisting of murine brain microcapillary endothelial cells and astrocytes (Fig. 1B,C). Further, 24-hour pre-treatment with NZ-8-061 did not modify diffusion of a low molecular weight fluorescent probe, the sodium fluorescein (Na-Fl), indicating that passage of NZ-8-061 was not a bystander effect due to its effects on BBB model permeability (Fig. 1D).

**Figure 1:**
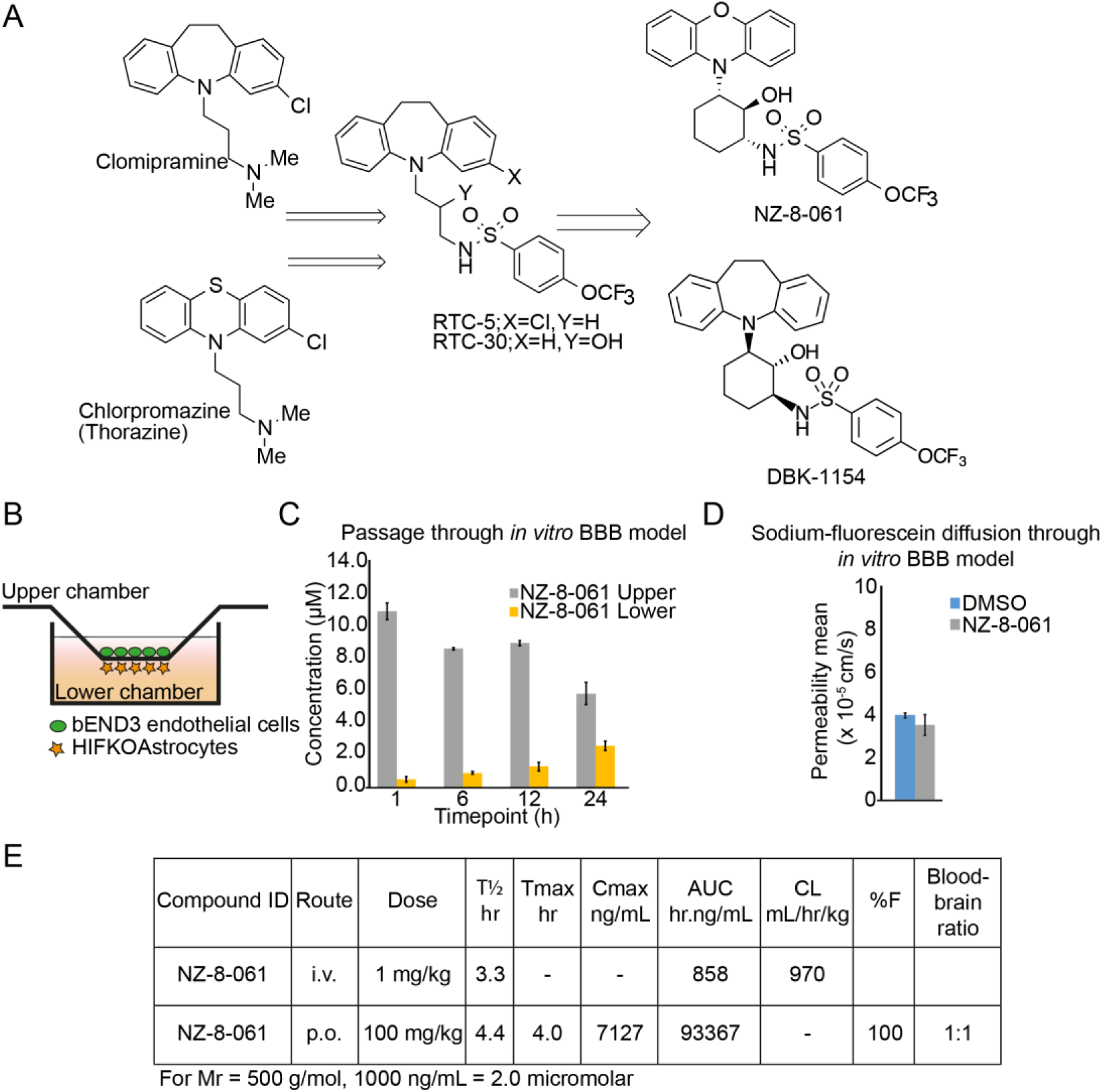
Blood-brain-barrier penetrance and mouse in vivo pharmacokinetics of NZ-8-061. **A)** An overview of development of small molecule activators of PP2A, NZ-8-061 and DBK-1154. The synthesis is described in detail in supplementary figure 1. **B)** Schematic presentation of in vitro BBB model which consists of murine endothelial cells and astrocytes as described in detail in (29). **C)** NZ-8-061 passage through the in vitro BBB model after addition of 15 µM dosage on the upper chamber at indicated timepoints. Data shown are means from two replicates ± SD. **D)** Sodium-fluorescein diffusion through the in vitro BBB after 24 hour of pretreatment with 15 µM NZ-8-061 on the upper chamber. Fluorescence signal of sodium-fluorescein was measured from lower chamber after 15 minutes. Data shown are means from two replicates ± SD. **E)** Mouse in vivo pharmacokinetic parameters (T½ hr, Tmax hr, Cmax ng/mL, AUC hr.ng/mL, CL mL/hr/kg, %F and blood-brain ratio) after 1 mg/kg or 100 mg/kg dosage via p.o. or i.v. of NZ-8-061.

To study brain penetration of NZ-8-061 *in vivo*, we performed a pharmacokinetic study by either administering 1 mg/kg i.v. or a bolus oral dose of 100 mg/kg. The *in vivo* pharmacokinetic parameters are shown in Figure 1E. NZ-8-061 shows 100% oral bioavailabilty based on dose adjusted fraction absorbed (%F) and moderate clearance as judged from half-life in plasma with T_1/2_ of 3 hr after I.V. dose. Peak plasma concentration after oral dose is around 14 micromolar and combined with moderate clearance and high AUC, shows significant, and sustained systemic exposure. Importantly, based on HPLC-MS/MS analysis from the whole brain homogenate, NZ-8-061 partitions into brain with a brain/plasma ratio of 1:1 at 6 hours post drug administration (Fig. 1E).

Together these results identify NZ-8-061 as an orally bioavailable, and BBB-permeable drug candidate for GB treatment.

### NZ-8-061 potently inhibits the viability of GB cells with heterogenous genetic background

Frequency of genetic mutations or deletions in any of the genes coding for core PP2A complex components in GB clinical isolates is negligible (Kaur *et al.*, 2016). However, we found that all studied GB cell lines expressed higher levels of PP2A inhibitor proteins (PAIPs); PME-1, CIP2A, SET, and ARPP-19 (Kauko and Westermarck, 2018), as compared to non-tumor fibroblasts (Fig. 2A). These results provide a non-genetic candidate mechanism for PP2A deregulation in GB cells.

**Figure 2:**
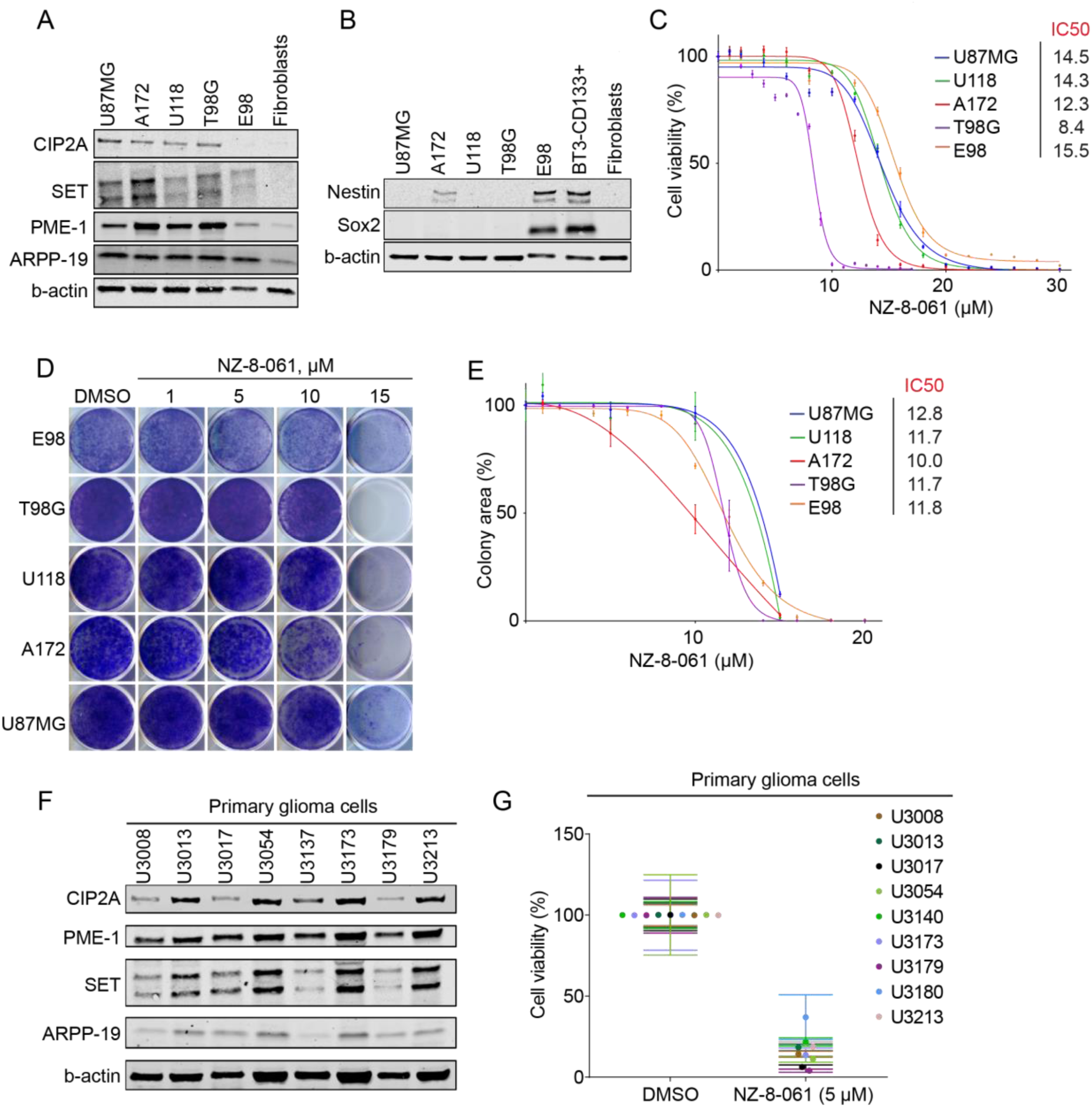
NZ-8-061 potently inhibits viability of molecularly heterogenous GB cells in vitro. **A)** Western blot of endogenous PP2A inhibitor proteins (ClP2A, SET, PME-1 and ARPP-19) and **B)** GB stem cell markers (Sox2 and Nestin) in indicated GB cell lines. BT3-CDl33+ (BT3 patient derived cells sorted for CD133) cells were used as a control for GB stem cell markers, and human fibroblasts were used as a negative control cell line in both. **C)** The dose-dependent effect of NZ-8-061 on the viability of indicated GB cell lines after 72 hours. Data shown are means from 6 replicates ± SD. lC5O values were calculated with GraphPad Prism 8. **D, E)** Colony growth reduction after NZ-8-061 treatment in GB cell lines with indicated concentrations. lC50 values for each cell line are shown. Data shown are means from 4 replicates ± SD. IC50 values were calculated with GraphPad Prism 8. **F)** Expression levels endogenous PPZA inhibitor proteins (CIPZA, SET, PME-1 and ARPP-19) in patient-derived primary glioma cells. **G)** Cell viability inhibition after NZ-8-061 (5 μM) treatment in patient-derived primary glioma cells. Data shown are means from 6 replicates ± SD.

Genomic heterogeneity and stemness characteristics are known to affect GB cell therapy responses. Genomic characteristics of the U87MG, A172, U118 and T98G cell lines were examined using the Cancer Cell Line Encyclopedia (CCLE) (Barretina *et al.*, 2012) database, whereas for the E98 cells published data was used (Claes *et al.*, 2008; Navis *et al.*, 2015). Collectively, the cell lines displayed a variety of known genomic alterations in GB (Supplementary Fig. 2A). However, the only common genomic change across all cell lines was copy number loss of CDKN2A (Supplementary Fig. 2A). Related to stemness properties, most of the established cell lines, except for E98, cultured with bovine serum supplements did not express any detectable levels of glioma stem cell markers SOX2 or Nestin (Fig. 2B).

We previously demonstrated notable kinase inhibitor resistance of T98G cells across different kinase families (Kaur *et al.*, 2016). To further examine kinase inhibitor responses of the used cell lines, we extracted the IC50 values for selected inhibitors from Genomics of Drug Sensitivity in Cancer database (https://www.cancerrxgene.org/). No information for E98 cell line was available. Consistently with their heterogenous mutation profiles (Supplementary Fig. 2A), each of the cell line have very variable sensitivity against different kinase inhibitors (Supplementary Fig. 2B). We experimentally validated Gefitinib resistance of three of the cell lines (Supplementary Fig. 2C).

These genetically diverse cell lines, displaying differential drug sensitivities, were then screened for their response to PP2A reactivation by NZ-8-061 in a cell viability assay. NZ-8-061 did induce a dose-dependent reduction in cell viability in all of the tested cell lines (Fig. 2C). Opposite to their kinase inhibitor responses, neither drastic differences, nor correlations of response with their genetic background, were found among the tested cell lines based on their IC50s. As a pharmacologic control, U87MG, U118, or A172 cells

were also treated with increasing doses of TRC-766, a structurally similar but biologically inactive derivate of SMAPs (Supplementary Fig. 3A)(Sangodkar *et al.*, 2017; McClinch *et al.*, 2018). While TRC-766 still binds PP2A, it is unable to reactivate PP2A even at a concentration of 20 µM *in vitro* (Sangodkar *et al.*, 2017). Notably, TRC-766 did not affect the viability of GB cells at concentrations up to 40 µM (Supplementary Fig. 3B). This strongly supports a dependence on PP2A reactivation for NZ-8-061-elicited killing of GB cells. Using a colony growth assay, NZ-8-061 treatment resulted in dose-dependent growth inhibition in all of the cell lines tested and in approximately the same concentration range as seen in the cell viability assay (Fig. 2D,E). Of note, micromolar dosing of SMAP is consistent with micromolar concentration of its molecular target PP2A in cancer cells, and required as serum binding of these compounds decreases their apparent potency in standard cell culture medium.

To extend these observations, we tested the therapeutic potential of NZ-8-061 in a series of thoroughly validated patient-derived primary glioma cells (Xie *et al.*, 2015). Notably, chosen cells represented all three molecular subtypes of GB according to Verhaak and colleagues (Wang *et al.*, 2017) (Supplementary Fig. 4A,B). Each of the patient-derived glioma cell line, cultured in serum-free neural stem cell media for the maintenance of stemness properties, expressed Nestin and Sox2, as well as PAIPs (Supplementary Fig. 4C and Fig. 2F). Further, they have a diverse genetic background (Supplementary Fig. 5A,B), and kinase inhibitor responses (Supplementary Fig. 6). All nine tested primary glioma cell lines displayed near to complete suppression of cell viability when treated with 5 µM NZ-8-061 (Fig. 2G). The lower working concentration of NZ-8-061 as compared to standard GB cell line cultures could be explained by the lack serum in the neural stem cell media, thus increasing the apparent potency of NZ-8-061.

Together these data demonstrate that SMAPs, as exemplified by NZ-8-061, have wide-spectrum therapeutic effect *in vitro* across human GB cells; regardless of their genetic background, disease subtype, or stemness properties.

### Preclinical activity of NZ-8-061 in an infiltrative intracranial GB model

To analyze *in vivo* therapeutic potential of oral dosing of NZ-8-061, we used an intracranial GB tumor model with luciferase-expressing/bioluminescent E98 cells (Claes *et al.*, 2008). Among the cell lines, E98 cells were selected as a model due to their stem-like properties (Fig. 2B), and infiltrative growth pattern in mouse brain recapitulating the human malignant glioma histology (Claes *et al.*, 2008) (Fig. 3A). Prior to the treatment, mice were randomized into two groups based on the tumor bioluminescence signal and using a protocol for optimized design and analysis of preclinical intervention studies *in vivo* (Laajala *et al.*, 2016). NZ-8-061 was thereafter orally dosed twice a day at 30 mg/kg. Based on pharmacokinetics and tissue distribution (Fig. 1E), the brain exposure of NZ-8-061 was estimated to transiently reach 10 µM, which is roughly comparable to the IC50 for colony formation assay with NZ-8-061 *in vitro* (Fig. 2E).

**Figure 3:**
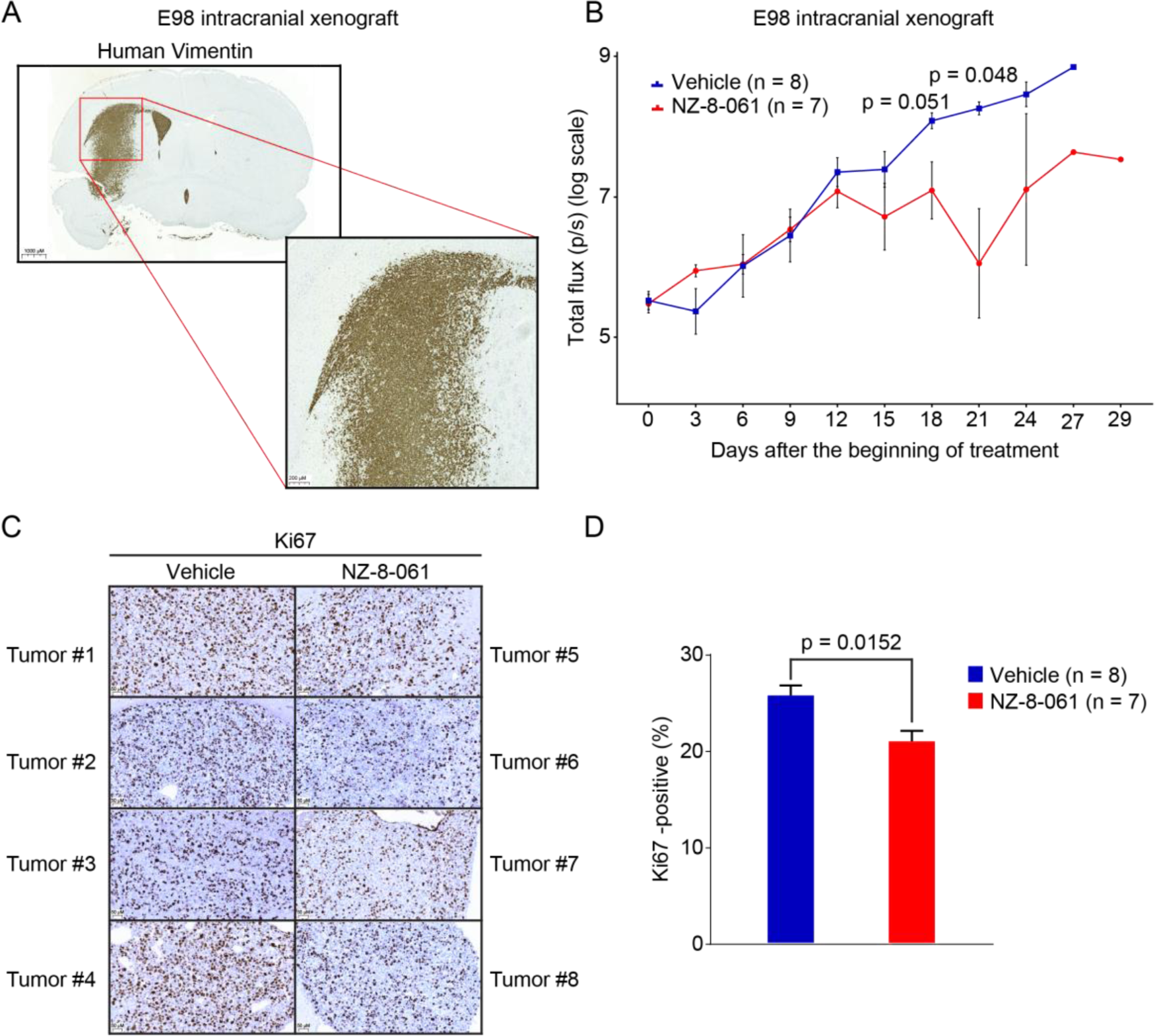
Therapeutic potential of oral dosing of NZ-8-061 as monotherapy in an infiltrative intracranial GB mouse model. **A)** Example picture of infiltrative growth of intracranial human E98-FM-Cherry cell line xenograft. The mouse brain tissue was stained with human specific vimentin antibody. **B)** Bioluminescence follow up of the intracranial human E98-FM-Cherry cell line xenograft growth during vehicle or 30 mg/kg NZ-8-061 treatment. When tumors were visible with bioluminescence, mice were randomized for either vehicle or NZ-8-061 groups. Data shown are means from 8 replicates ± SEM, p-values by Student’s t-test. **C)** Representative images of Ki-67 staining from 4 vehicle and 4 NZ-8-061-treated end-point tumors from B). **D)** Quantification of Ki-67 positivity from C). Data shown are means of % of Ki-67 positive tumor cells from 7-8 tumors/group ± SD, *, p-value by Mann-Whitney test.

Intracranial tumor growth was followed by bioluminescence measurements every third day. Notably, NZ-8-061 induced tumor growth stasis at day 12 and displayed a significant reduction in tumor size at later timepoints (Fig. 3B). The therapeutic response was validated by significant inhibition in tumor cell Ki67 expression in the end-point tumors when comparing treated and non-treated groups (Fig. 3C, D).

### DBK-1154, with higher degree of brain/blood distribution, and more efficient *in vitro* activity, increases survival of mice bearing orthotopic GB tumors

Despite its significant *in vivo* efficacy in reducing intracranial tumor growth with roughly IC50 dosing (Fig. 3B-D), NZ-8-061 monotherapy failed to improve the mouse survival (Supplementary figure 7A). We therefore next tested the BBB permeability and GB cell killing properties of DBK-1154. DBK-1154, with a dibenzoazepine tricyclic, has a hydrocarbon bridge versus an oxygen bridge in NZ-8-061 (Fig. 1A), making DBK-1154 somewhat more lipophilic. The calculated LogP (log octanol-water partition coefficient), a measure of lipophilicity, is higher for DBK-1154 (cLogP 7.0) than NZ-8-061 (cLogP 6.6). Additionally, tPSA (total polar surface area) is higher for NZ-8-061 versus DBK-1154 (88 vs 79 A^2^). Lower tPSA and higher cLogP generally correlate with higher CNS distribution (Kelder *et al.*, 1999).

*In vivo* pharmacokinetic parameters for DBK-1154 in mouse are shown in figure 4A and in supplementary figure 7B. DBK-1154 is orally bioavailable, however it showed significantly lower systemic plasma exposure compared to NZ-8-061. Importantly, in addition to *in* vitro BBB permeability (Supplementary Fig. 8A), the *in vivo* evaluation of distribution of DBK-1154 into the CNS showed a 2.3-fold higher concentration in brain tissue versus plasma (Fig. 4A).

**Figure 4:**
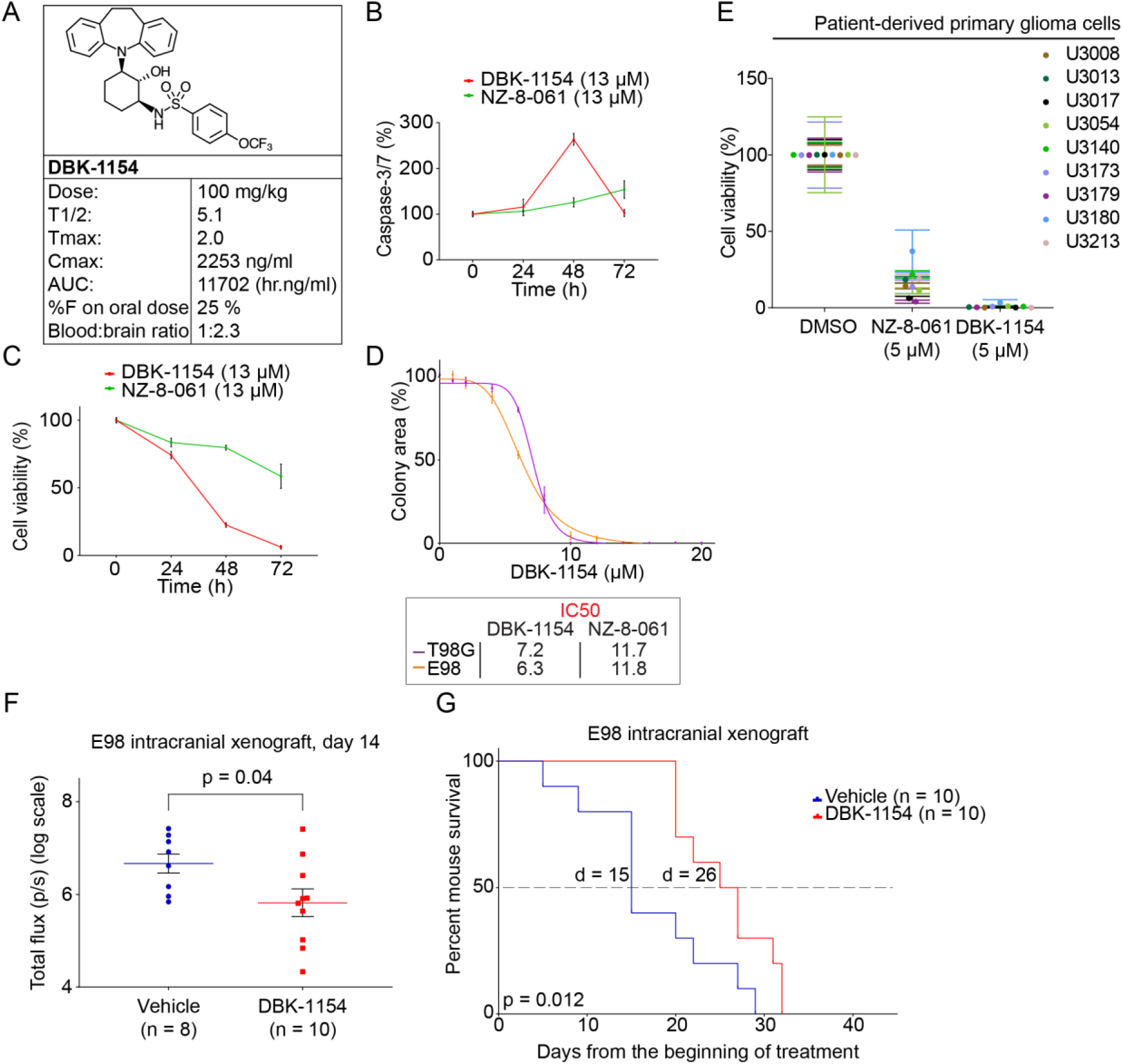
Oral dosing of DBK-1154 increases survival of mouse with intracranial infiltrative GB xenograft. **A)** Mouse in vivo pharmacokinetic parameters (T½ hr, Tmax hr, Cmax ng/mL, AUC hr.ng/mL, CL mL/hr/kg, %F and blood-brain ratio) after 100 mg/kg dosage via p.o. of DBK-1154. **B)** Time-dependent Caspase -3/7 activation in E98 cells, after either DT-061 or DBK-1154 treatment (13 µM). **C)** Parallel with the caspase 3/7 assay, a cell viability assay was run with same timepoints and concentration. Data shown are means from 6 replicates ± SD. **D)** Colony growth reduction after DBK-I l 54 treatment in GB cell lines in indicated concentrations. ICSO comparison between DBK-1 154 and NZ-8-061 is shown in the insert. Data shown are means from 4 replicates ± SD. **E)** Cell viability inhibition after DBK-1154 (5 μM) or NZ-8-061 (5 μM) treatment in patient-derived primary glioma cells cultured in serum-free neural stem cell medium. Data shown are means from 6 replicates ± SD. **F)** Bioluminescence comparison on the day 14 during the orthotopic E98 in vivo model between vehicle or 100 mg/kg DBK-1154 treatment. Data shown are means from 8 vehicle and 10 DBK-1154 treated mice ± SEM, *, p-value by Student’s t-test. **G)** Survival of mice with orthotopic E98 xenografts after vehicle or 100 mg/kg DBK-1154 treatment, *, P < 0.05 by Gehan-Breslow-Wilcoxon test. Mice were randomized to 2 groups of I0 mice each based on bioluminescence signal before starting the treatment. Median survival was increased with DBK-l I54 treatment from 15 days to 26 days.

Based on the IC50-value, higher potential of DBK-1154 was seen in a long-term colony growth assay in which the IC50 for DBK-1154 was almost two-fold lower than that of NZ-8-061 in both cell lines (Fig. 4D, insert). In addition, DBK-1154 was qualitatively different from NZ-8-061 in terms of its apoptosis inducing potential. With single 13 µM dosing, only DBK-1154 induced Caspase-3/7 cleavage at 48 hours (Fig. 4B). This was reflected in a significant time-dependent difference in inhibition of cell viability also starting at 48 hours (Fig. 4C). The higher potency of DBK-1154, when compared to NZ-8-061, was also seen across patient-derived primary glioma cells in which DBK-1154 induced complete inhibition of cell viability (Fig. 4E). Importantly, using the same cell culture conditions, most of the tested kinase inhibitors did not reach even 50% reduction in cell viability with up to 10 µM concentration in all five of the tested patient-derived primary glioma cells (Supplementary Fig. 6).

Next, we proceeded to test DBK-1154 in the same intracranial E98 model as with NZ-8-061. To compensate for the lower oral bioavailability of DBK-1154, it was dosed at 100 mg/kg, twice daily, using the same homogeneous formulation, with expected transient brain exposure of about 8 micromolar twice a day. This level of CNS exposure is roughly comparable to the IC50 for colony formation for DBK-1154 *in vitro* (Fig. 4D). DBK-1154 therapy caused tumor growth reduction after day 5 (Supplementary Fig. 8D), and reaching statistically significant difference at day 14 (Fig. 4F). Further, by using a linear mixed effects model that pooled the control groups from both experiments with the assumption that only the starting luciferase signals differ, the effects with DBK-1154 on linear tumor growth were found to be highly significant (p=0.00316, Bonferroni p=0.00632). Further, while NZ-8-061 monotherapy did not increase the survival of the intracranial tumor bearing mice (Supplementary figure 7A), the higher *in vitro* efficiency and *in vivo* brain penetrance of DBK-1154 translated to a significant, almost 2-fold longer overall mouse survival. The median survival was 15 days for the control group, and 26 days for the DBK-1154 treated group (p=0.012) (Fig. 4G). Together these data indicate that of the studied SMAPs, DBK-1154 has a greater potential as a GB therapy candidate molecule.

Considering the potential future development of DBK-1154 derivatives for GB therapy, DBK-1154 was evaluated in an acute rat pilot (non-GLP) toxicology study at doses up to 800 mg/kg daily. Consistently with previous SMAP *in vivo* studies (Sangodkar *et al.*, 2017; Kauko *et al.*, 2018; McClinch *et al.*, 2018), no significant body weight loss, deaths, or adverse behavioral or neurological effects were observed. The major morphologic finding was hepatocellular hypertrophy (panlobular) in all test article-treated groups, with increased severity as the dose level increased. This observation was likely related to compound specific pregnane X receptor (PXR) agonist activity, and was considered an adaptive rather than toxic effect. These findings indicate for a clear therapeutic window between normal and cancer cells *in vivo.*

Collectively these results provide proof-of-principle evidence for preclinical *in vivo* efficacy, and acceptable safety profile of SMAPs as novel candidate class of BBB penetrable, tumor suppressor reactivation therapeutics. Notably, the efficacy of SMAPs did not depend on the subtype of GB, or the genomic alterations, in either established GB cell lines or in patient derived primary glioma cells. Moreover, SMAPs were found to be superior to a range of kinase inhibitors in their capacity to kill patient derived primary glioma cells. These results provide the first indications that PP2A reactivation might be able to challenge the current paradigm in GB therapies which has been strongly focused on targeting specific genetically altered cancer drivers with highly specific inhibitors (Verhaak *et al.*, 2010; Barretina *et al.*, 2012; Brennan *et al.*, 2013). PP2A is known to simultaneously target a number of cancer driver pathways and pro-apoptotic mechanisms (Perrotti and Neviani, 2013; Kauko and Westermarck, 2018). We envision that these wide-spectrum effects may explain the very robust preclinical survival effects across kinase inhibitor resistant cell lines harboring heterogenous genetic drivers.

Another serious obstacle for development of GB therapies is inadequate exposure of most developed molecules in the CNS due to the BBB (Harder *et al.*, 2018). Here we show that both SMAPs have retained the BBB penetration properties of tricyclics from which they were derived. This was not obvious as SMAPs contain a significant modification of the original tricyclics with loss of a polar amine salt and addition of sulfonamide and hydroxyl moieties. The observation that out of the studied SMAPs, DBK-1154 preferably partitions into the brain, can be explained by its lower tPSA and higher cLopP. Consequently, this better brain partitioning can partly explain the significant almost 2-fold prolongation of survival of mice with DBK-1154 versus NZ-8-061. In addition to GB, poor small molecule BBB penetration is a serious problem also for the therapy of brain metastasis from many non-CNS cancer types. As these are more common than primary brain tumors such as GB, the presented proof-of-principle data for usefulness of SMAPs as CNS therapeutics might be relevant for much larger patient population than only GB patients.

In addition to their different *in vivo* brain penetrance properties, NZ-8-061 seems to have cytostatic effects at the doses tested, whereas DBK-1154 most probably induces both, decreased proliferation and cell death. On the other hand, a SMAP derivative DBK-766 defective in PP2A reactivation (Sangodkar *et al.*, 2017; McClinch *et al.*, 2018), failed to suppress GB cell viability. The intracellular pathways involved in the apoptosis induction by DBK-1154 *in vitro* were not addressed in this concise communication, but clearly remains as an important future question to be addressed. As such, the results provide a clear indication that chemical structure of DBK-1154 could serve as a scaffold for further development of even more potent small molecule activators of PP2A for GB therapy. Following this rationale, a medicinal chemistry program identifying DBK-1154 analogs with improved oral bioavailability is currently ongoing.

PP2A activity is known to modulate kinase inhibitor responses in hematological and solid cancers (Neviani *et al.*, 2013; Kaur *et al.*, 2016; Kauko *et al.*, 2018). *In vivo*, combination studies with SMAPs demonstrated significant tumor regression in KRAS-mutant lung cancer xenograft model with MEK inhibitors (Kauko *et al.*, 2018). Therefore, the results reported here not only serve as a proof-of-principle for the feasibility of SMAPs as novel glioblastoma therapeutics, but also pave the way for future combination studies with kinase inhibitors, and potentially also other type of therapies in brain cancers. As PP2A inhibition is also one of the pathogenic mechanisms in Alzheimer’s disease, our data demonstrating BBB penetration and *in vivo* therapeutic effects of SMAPs in intracranial model might also serve as a landmark for use of SMAPs also in other CNS pathologies.

## Methods

Further information is provided in Supplemental Methods.

## Data availability statement

Data is available from the corresponding author Jukka Westermarck upon request.

