## Supplementary data for "Monotherapy efficacy of BBB-permeable small molecule activators of PP2A in glioblastoma" (from another version of this preprint: version 1, 2019-09-23)

Supplementary Figure 1.

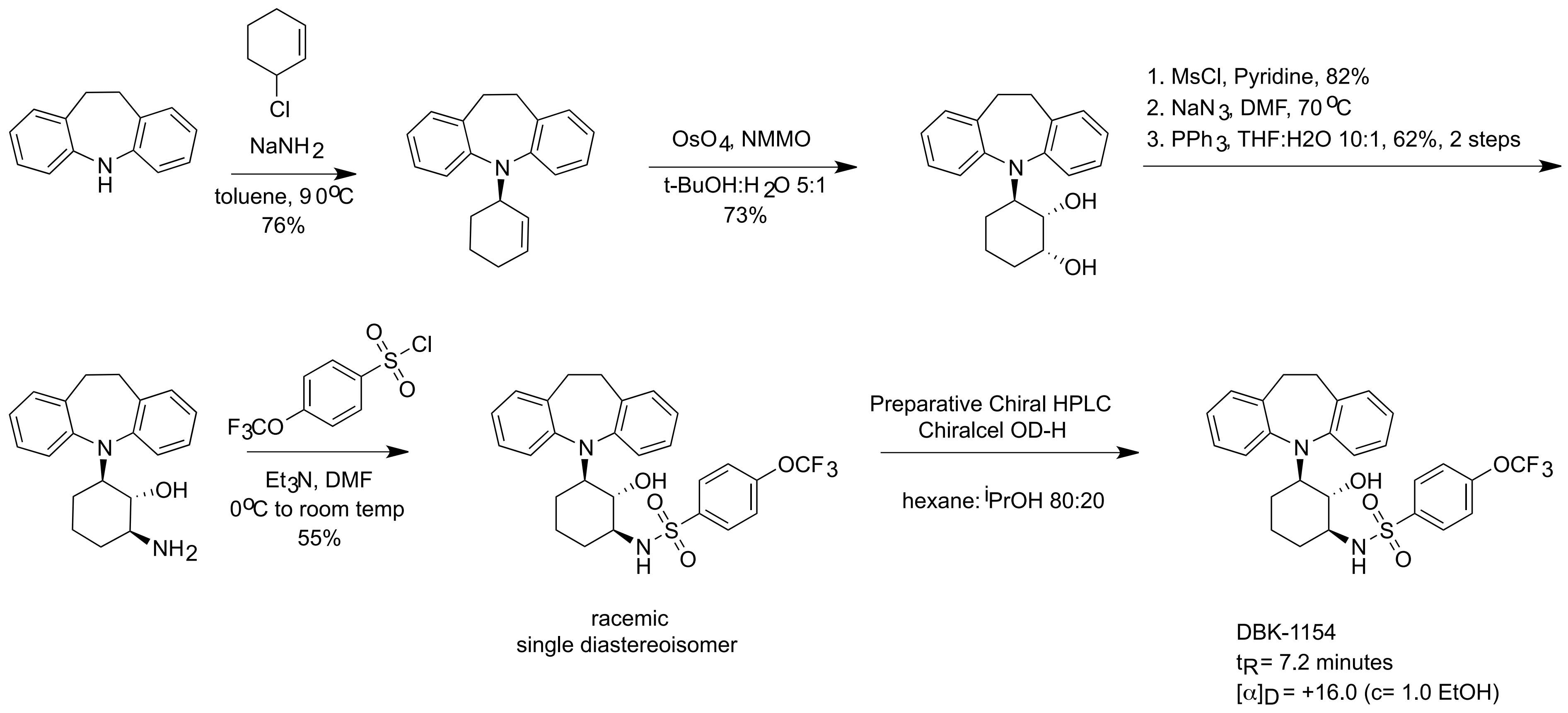

Supplementary Figure 2.

A

### Genetic alterations in GBM cell lines

| Common genetic alterations | U87MG | A172 | U118 | T98G | E98 |
|---|---|---|---|---|---|
| RTK | | | | | |
| EGFR | CV gain | Coding silent | | | |
| PDGFRA | | | Missense | | |
| PDGFRB | | Over expression | | | |
| MET | | | | | Mutation |
| PI3K | | | | | |
| PIK3CB | | Missense | | | |
| PIK3C2A | Over expression | | | | |
| PTEN | Mutation | Loss of CN | Mutation | Missense | |
| MAPK | | | | | |
| NF1 | Deletion | | | | |
| BRAF | Over expression | | | | |
| KRAS | Under expression | | | | |
| NRAS | Over expression | | | | |
| P53 | | | | | |
| TP53 | | | Missense | Missense | |
| RB1 | | | | | |
| CDKN2A | Loss of CN | Loss of CN | Loss of CN | Loss of CN | Loss of CN |
| CDKN2B | Loss of CN | Loss of CN | Loss of CN | | |
| CDKN2C | Loss of CN | | | Loss of CN | |
| CCND2 | | | Missense | | |
| RB1 | | Missense | | | |
| Chromatin modifier | | | | | |
| ATRX | Missense | | | | |
| Other | | | | | |
| MGMT | +++ | | | | |

B

| | | IC50 (μM) | | | |
|---|---|---|---|---|---|
| | | U87MG | T98G | A172 | U118 |
| EGFR | Gefinitib | **44.84** | 20.32 | 17.63 | 33.00 |
| | Osimertinib | **15.13** | 12.35 | 4.14 | 5.33 |
| VEGFR | Sorafenib | **35.93** | 16.33 | 7.22 | 11.79 |
| | Cediranib | 7.27 | **14.15** | 1.71 | 8.28 |
| MEK | Trametinib | 2.72 | **10.26** | 1.50 | 2.88 |
| MET | Foretinib | **2.65** | 2.42 | 1.33 | 2.02 |
| | Savolitinib | **13.74** | 13.51 | 6.84 | 3.92 |
| BRAF | Dabrafenib | **381.38** | 161.77 | 52.09 | 341.55 |
| mTOR | Rapamycin | 0.05 | 0.05 | 0.04 | **0.07** |
| PI3K | Alpelisib | 62.61 | 45.46 | 119.21 | **399.87** |
| | Buparlisib | 3.00 | **3.25** | 1.23 | 2.51 |
| CDK 4/6 | Palbociclib | 58.96 | **62.88** | 18.80 | 52.25 |
| AKT | Uprosertib | **90.44** | 23.86 | 15.72 | 26.29 |
| | MK-2206 | 28.86 | **60.39** | 46.15 | 27.82 |

C

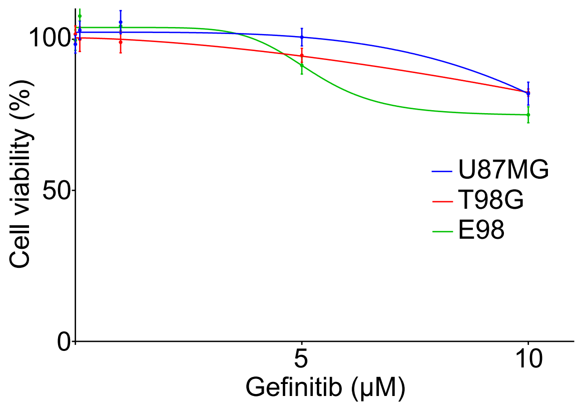

Supplementary Figure 3.

A

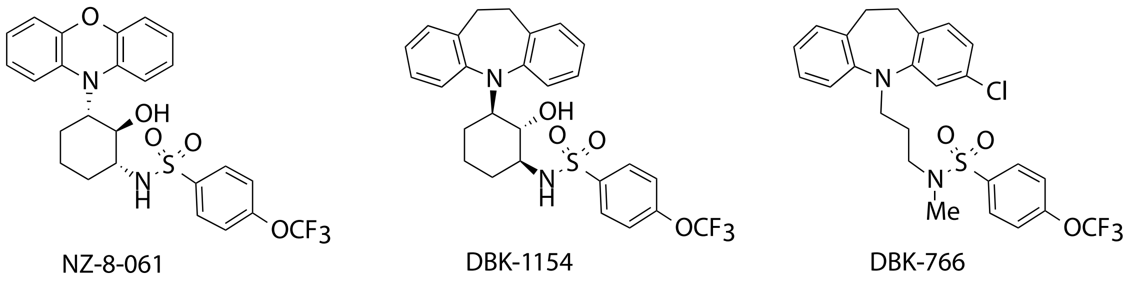


B

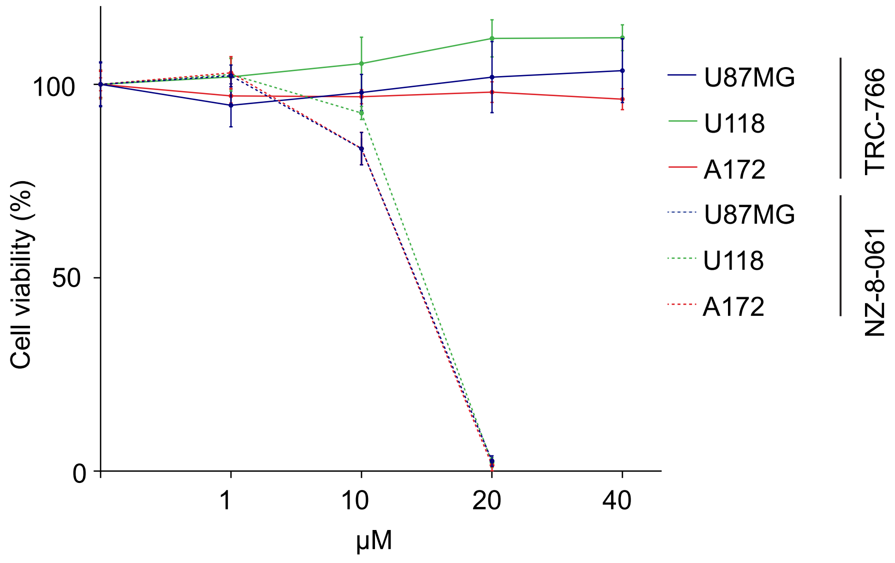

Supplementary Figure 4.

A

| | P value of cell line for subtypes | | | | | | | | |
|---|---|---|---|---|---|---|---|---|---|
| Subtype | U3008 | U3013 | U3017 | U3054 | U3173 | U3180 | U3137 | U3213 | U3179 |
| 'Proneural_pval' | 1 | 0.003 | 0.996 | 0.052 | 0.101 | 0.999 | 0.728 | 0.698 | 0.718 |
| 'Classical_pval' | 0.106 | 0.996 | 0.006 | 1 | 0.946 | 0.001 | 0.258 | 0.543 | 0.462 |
| 'Mesenchymal_pval' | 0.783 | 0.970 | 0.952 | 0.097 | 0.023 | 0.996 | 0.029 | 0.002 | 0.996 |

B

Subtypes of primary glioma cell lines

| | |
|---|---|
| U3008 | Classical |
| U3013 | Proneural |
| U3017 | Classical |
| U3054 | Proneural |
| U3140 | Proneural |
| U3173 | Mesenchymal |
| U3179 | Classical |
| U3180 | Classical |
| U3213 | Mesenchymal |

C

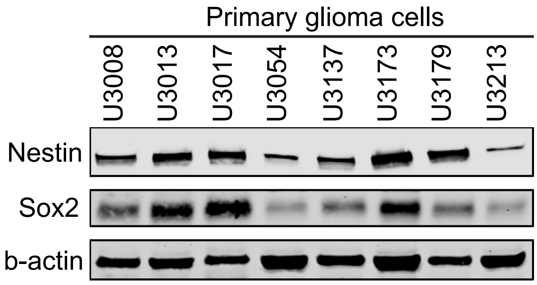

Supplementary Figure 5.

A

Copy number alterations in patient-derived glioma stem cell lines

| | U3008 | U3013 | U3017 | U3054 | U3140 | U3173 | U3179 | U3180 | U3213 |
|---|---|---|---|---|---|---|---|---|---|
| 'ATRX' | 0.042 | 0.306 | -0.534 | 0.082 | 0.063 | -0.13 | -0.565 | -0.634 | 0.021 |
| 'BRAF' | 0.386 | 0.356 | 0.405 | 0.413 | 0.358 | 0.376 | 0.415 | 0.398 | 0.408 |
| 'CCND2' | 0.047 | 0.244 | 0.065 | 0.077 | 0.076 | 0.009 | 0.062 | 0.039 | 0.039 |
| 'CDKN2A' | -1.996 | 0.093 | -1.3 | -0.309 | -1.835 | -0.03 | -1.853 | -1.97 | -2.162 |
| 'CDKN2B' | -1.996 | 0.093 | -1.3 | -0.309 | -1.835 | -0.03 | -1.853 | -1.97 | -2.162 |
| 'CDKN2C' | 0.041 | -0.142 | -0.457 | 0.024 | -0.132 | 0.043 | 0.052 | 0.033 | -0.106 |
| 'EGFR' | 0.38 | 0.345 | 0.396 | 0.522 | 0.3 | 0.34 | 0.638 | 0.434 | 0.407 |
| 'KRAS' | 0.068 | 0.275 | 0.065 | 0.13 | 0.058 | -0.009 | 0.038 | 0.009 | 0.039 |
| 'MET' | 0.403 | 0.356 | 0.392 | 0.61 | 0.437 | 0.299 | 0.294 | 0.32 | 0.408 |
| 'MGMT' | -0.521 | -0.146 | -0.482 | -0.328 | -0.535 | -0.279 | -0.521 | 0.036 | -0.434 |
| 'NF1' | 0.046 | 0.236 | 0.075 | 0.204 | 0.052 | -0.065 | -1.97 | 0.073 | 0.037 |
| 'NRAS' | 0.061 | 0.052 | 0.066 | 0.026 | -0.185 | 0.186 | 0.046 | -0.001 | -0.106 |
| 'PDGFRB' | 0.06 | 0.043 | 0.05 | -0.095 | 0.049 | -0.026 | 0.043 | 0.019 | 0.022 |
| 'PIK3C2A' | 0.041 | 0 | 0.067 | 0.017 | 0.008 | -0.064 | 0.05 | 0.013 | 0.046 |
| 'PIK3CB' | 0.03 | 0.109 | 0.057 | -0.075 | 0.072 | 0.139 | 0.046 | -0.015 | -0.082 |
| 'PTEN' | -0.326 | 0.001 | -0.482 | -1.735 | -1.73 | -1.737 | -0.369 | -0.407 | -0.434 |
| 'RB1' | 0.033 | -0.39 | -0.465 | -0.052 | -0.167 | -0.411 | 0.035 | 0.016 | 0.035 |
| 'TP53' | 0.041 | 0.236 | 0.043 | -0.378 | 0.034 | -0.026 | 0.046 | 0.61 | 0.002 |

B

Point mutations in patient-derived glioma stem cell lines

| Gene name | U3008 | U3013 | U3017 | U3054 | U3140 | U3173 | U3179 |
|---|---|---|---|---|---|---|---|
| 'ATRX' | 0 | 0 | 0 | 0 | 0 | 0 | 0 |
| 'BRAF' | 0 | 0 | 0 | 0 | 0 | 0 | 0 |
| 'CCND2' | 0 | 0 | 0 | 0 | 0 | 0 | 0 |
| 'CDKN2A' | 0 | 0 | 0 | 0 | 0 | 0 | 0 |
| CDKN2B' | 0 | 0 | 0 | 0 | 0 | 0 | 0 |
| 'CDKN2C' | 0 | 0 | 0 | 0 | 0 | 0 | 0 |
| 'EGFR' | 0 | 0 | 0 | 0 | 0 | 0 | 0 |
| 'KRAS' | 0 | 0 | 0 | 0 | 0 | 0 | 0 |
| 'MET' | 0 | 0 | 0 | 0 | 0 | 0 | 0 |
| 'MGMT' | 0 | 0 | 0 | 0 | 0 | 0 | 0 |
| 'NF1' | 0 | 0 | 0 | 0 | 0 | 0 | 0 |
| 'NRAS' | 0 | 0 | 0 | 0 | 0 | 0 | 0 |
| 'PDGFRB' | 0 | 0 | 0 | 0 | 0 | 0 | 0 |
| 'PIK3C2A' | 0 | 0 | 0 | 0 | 0 | 0 | 0 |
| PIK3CB' | 0 | 0 | 0 | 0 | 0 | 0 | 0 |
| 'PTEN' | 0 | 0 | 0 | 0 | 0 | 0 | 0 |
| 'RB1' | 0 | 0 | 0 | 0 | 0 | 1 | 0 |
| TP53' | 0 | 1 | 0 | 1 | 0 | 1 | 0 |

Supplementary Figure 6.

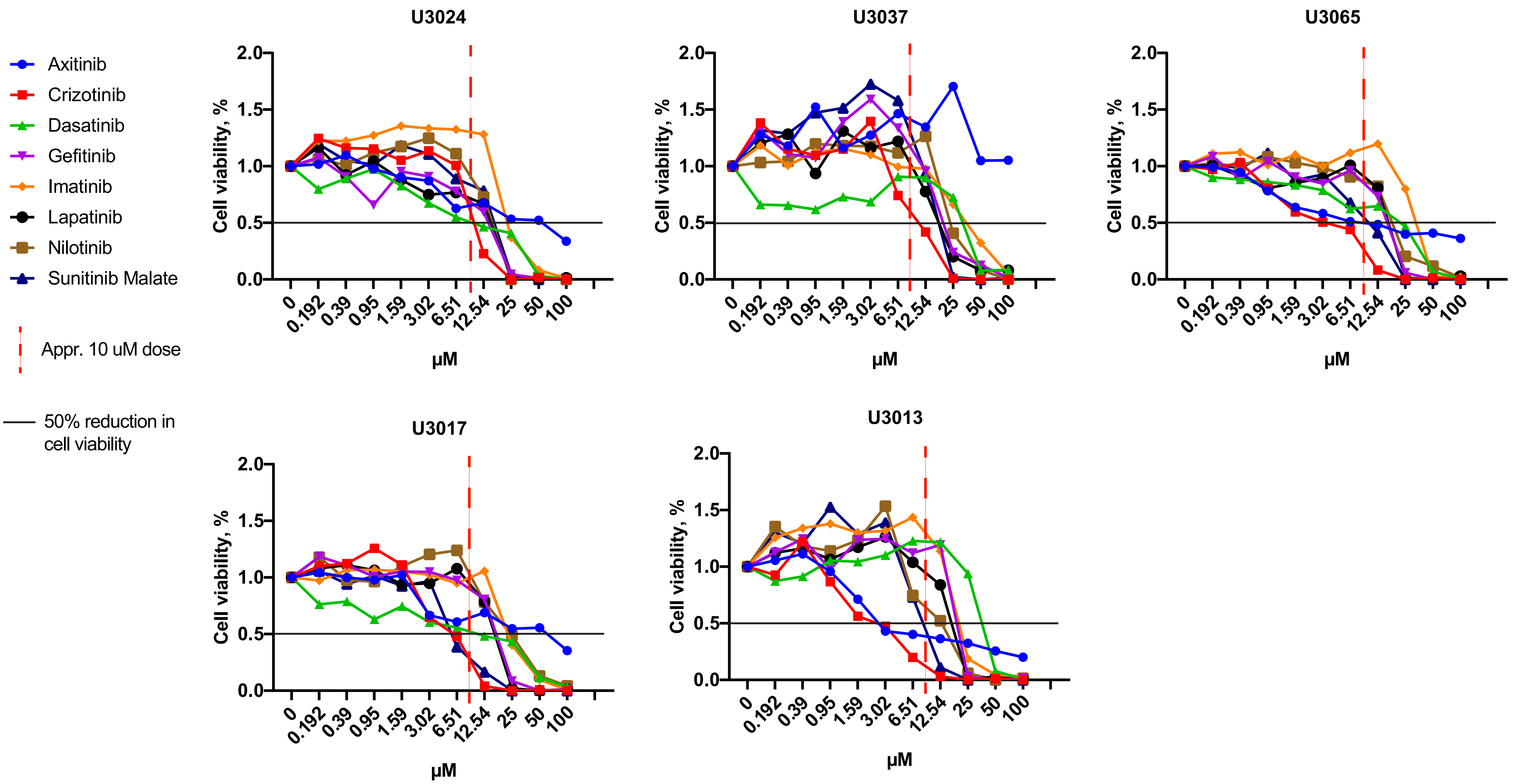

Supplementary Figure 7.

A

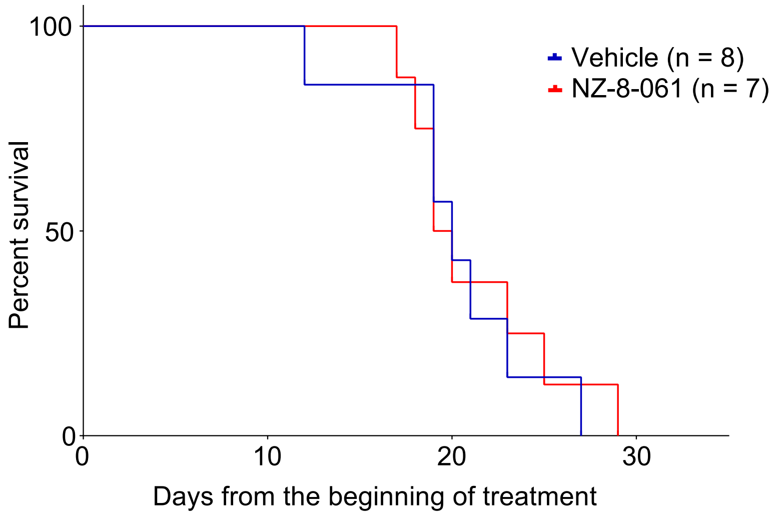


B

| Compound ID | Route | Dose | T½ hr | Tmax hr | Cmax ng/mL | AUC hr.ng/mL | CL mL/hr/kg | %F | Blood-brain ratio |
|---|---|---|---|---|---|---|---|---|---|
| DBK-1154 | i.v. | 1 mg/kg | 2.1 | - | 1478 | 495 | 1950 | | |
| DBK-1154 | p.o. (soln.) | 100 mg/kg | 5.1 | 2.0 | 2253 | 11702 | - | 23 | 1:1 |
| DBK-1154 | p.o. (susp.) | 100 mg/kg | 3.6 | 4.0 | 1200 | 7130 | - | 14 | 1:1 |

For Mr = 500 g/mol, 1000 ng/mL = 2.0 micromolar

Supplementary Figure 8.

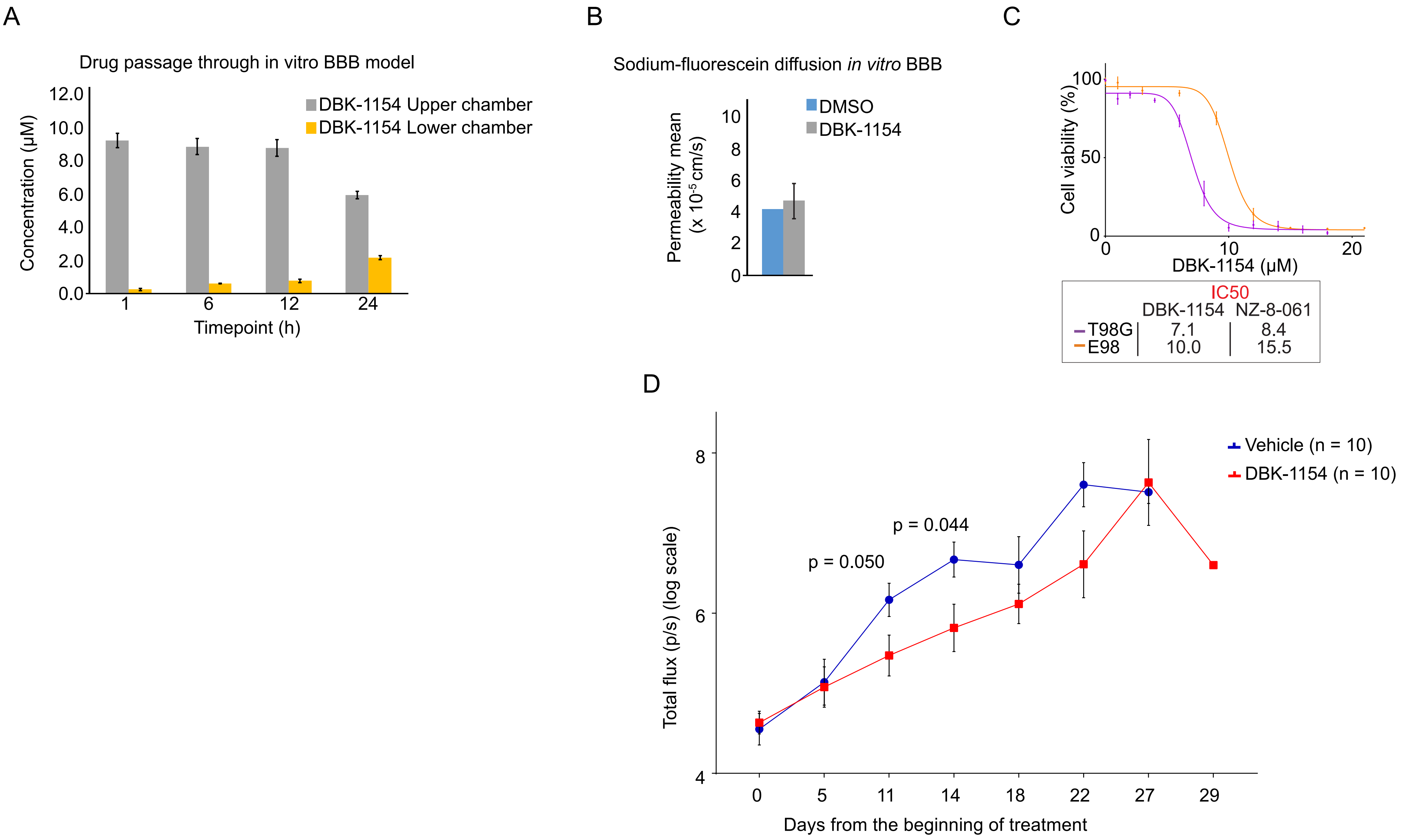
